# A Prediction Method of Land Use Type Evolution Based on Long and Short-Term Memory Networks—Taking Gansu Province as an Example

**DOI:** 10.1101/2024.12.20.629836

**Authors:** Shiqi Zhang, Chuhui Cao

## Abstract

In the context of escalating global climate change and human activities, understanding the driving mechanisms behind land use change and predicting future trends is crucial. This study takes Gansu Province as a case, using land use type data from 1990 to 2020 to construct a Long Short-Term Memory (LSTM) model to predict land use changes over the next decade (2021–2030). The results indicate that land use types in Gansu Province exhibit significant dynamic changes, with forest area continuously increasing, built-up land rapidly expanding, and areas of unused land and grassland significantly decreasing. These changes reflect the combined effects of ecological protection policies, urbanization, and land development. The model’s predictions suggest that built-up land has absorbed substantial areas of unused land, grassland, and cultivated land, with accelerated urbanization. Forest area growth is attributed to the implementation of ecological restoration policies, while grassland and water areas show fluctuating changes, and the area of unused land continues to decrease. The findings not only provide data support for land resource management and ecological protection, but also offer scientific evidence for the formulation of sustainable land use policies, which can serve as an important reference for the sustainable use and management of land resources in Gansu Province and similar regions.

## 1 Introduction

Land use change is a complex and dynamic process driven by both natural factors and human activities, with profound implications for the balance of ecosystems, regional development directions, and socioeconomic transformation. In the context of escalating global climate change and intensifying human activities, understanding the driving mechanisms behind land use change and predicting future trends is not only a frontier issue in scientific research, but also a key to achieving effective land resource management, promoting sustainable development, and fostering ecological restoration [1-3]. Through in-depth research on land use change, valuable scientific evidence can be provided for resource allocation optimization, the formulation of ecological protection policies, and regional economic planning [4,5].

Gansu Province, located in the northwest of China, is characterized by a complex geographical environment and a fragile ecosystem. Its land use types are diverse, including arable land, forestland, grassland, water bodies, unused land, and built-up land. The dynamic changes among these land types directly reflect the combined influence of natural conditions and human activities [6,7]. In recent years, significant changes in land use patterns have occurred in Gansu due to the acceleration of urbanization, the gradual implementation of ecological protection policies, and the growing demands of agricultural development. For example, forestland has steadily increased under the promotion of ecological protection policies, while grassland and unused land have gradually decreased due to economic development needs [8]. These changes not only involve the allocation of natural resources but also profoundly impact the adjustment of regional ecological functions and economic structures [9-11]. Therefore, forecasting future land use changes in Gansu Province is of great significance for addressing issues such as ecological degradation, water resource management, and regional sustainable planning.

Despite significant advancements in land use change modeling, traditional methods still face limitations when addressing the complexity of dynamic land use changes. Conventional statistical models, such as linear regression and Autoregressive Integrated Moving Average (ARIMA) models, have been widely used to analyze the temporal patterns of land use change. However, these methods often rely on linear assumptions and exhibit considerable limitations when dealing with nonlinear and complex dynamic changes [12]. For example, researchers found that when applying the ARIMA model to analyze changes in arable land, it produced significant errors in handling nonlinear trends and struggled to accurately capture the complex dynamic relationships underlying land use changes [13]. To overcome the shortcomings of traditional methods, machine learning techniques have gradually been introduced into land use change modeling [14-16]. Models such as Support Vector Machine (SVM) and Random Forest have demonstrated strong performance in identifying the driving factors of land use change and in remote sensing data classification. However, these models have relatively limited capabilities in modeling temporal dependencies, making it difficult to capture potential trends in long-term changes [17-19]. This limitation has driven researchers to explore more powerful modeling tools to conduct a more comprehensive analysis of dynamic land use changes [20].

In recent years, deep learning models, particularly Long Short-Term Memory (LSTM) networks, have demonstrated exceptional capabilities in the field of time series analysis. Through their unique memory units and forget mechanisms, LSTM networks are able to capture the nonlinear characteristics of long-term dependencies within data [21]. These models have been widely applied in fields such as climate modeling, hydrological forecasting, and land use change. For instance, researchers used the LSTM model to predict urban expansion in the Yangtze River Delta region, successfully capturing the temporal dynamic features of land use change and significantly improving the prediction accuracy [22]. Additionally, researchers verified the effectiveness of the LSTM model in handling complex temporal relationships while studying forest cover change in Northeast China, further demonstrating its potential in land use change prediction [23,24].

Although the LSTM model has demonstrated significant advantages in land use change modeling, its application remains relatively limited under diverse ecological and socio-economic conditions. Gansu Province, as an ecologically fragile region with complex land use types, provides an ideal study area to explore the applicability of the LSTM model. This study aims to develop an LSTM-based time series forecasting model to comprehensively analyze the land use change history of Gansu Province from 1990 to 2020 and to predict land use changes for the period 2021-2030. The study will focus on assessing the dynamic patterns of different land use types, exploring the mechanisms of key driving factors, and further analyzing the implications of the prediction results for regional planning and ecological protection. The significance of this study lies not only in enhancing the predictive capability of land use changes in Gansu Province, but also in providing practical support for addressing pressing issues such as land degradation, urban expansion, and water resource management. By forecasting future land use trends, the study can help identify potential areas at risk of ecological degradation and provide scientific evidence for policy interventions.

## 2 Materials and methods

### 2.1 Overview of the study area

Gansu Province, located in the northwest of China, lies between longitudes 92°13′ to 108°46′ east and latitudes 32°11′ to 42°57′ north, covering a total area of approximately 425,900 square kilometers, accounting for 4.4% of the national land area. The province features a topography characterized by a high west and low east gradient, with a distinct stepped transition. It serves as the intersection of the Tibetan Plateau, the Loess Plateau, and the Inner Mongolia Plateau within China’s natural geographic structure. The region’s terrain is diverse, including mountains, hills, plateaus, basins, and plains. The highest elevation in the province is approximately 5,547 meters, while the lowest is around 500 meters, resulting in a significant elevation drop and a relatively fragile ecological environment. Gansu Province is rich in land resources, with diverse land use types, primarily including arable land, forestland, grassland, unused land, and built-up land. The province’s arable land is mainly concentrated in the Hexi Corridor and the Loess Plateau region, which are key agricultural production areas. Forests are predominantly located in the southern mountainous areas, where the implementation of ecological protection policies has gradually increased the forest coverage rate. Grasslands are widely distributed in the arid and semi-arid regions in the northwest of the province, serving as an important foundation for livestock development. However, in recent years, grassland degradation has become a significant issue due to climate change and human activities. Unused land is primarily found in the Gobi Desert and desert areas, where there is considerable potential for land resource development. Climatically, Gansu Province falls within the temperate monsoon climate zone and the temperate arid-semiarid climate zone, characterized by significant precipitation variation. The southeastern part is humid and receives abundant rainfall, while the northwestern part is arid and receives little rainfall. Water resources are unevenly distributed, and the ecological environment is fragile. In recent years, with the advancement of ecological protection projects (such as the Three-North Shelter Forest Program and the Grain-for-Green Program), the ecological environment of Gansu has gradually improved. However, issues such as soil erosion, desertification, and grassland degradation remain prominent. The specific study area is shown in Figure 1.

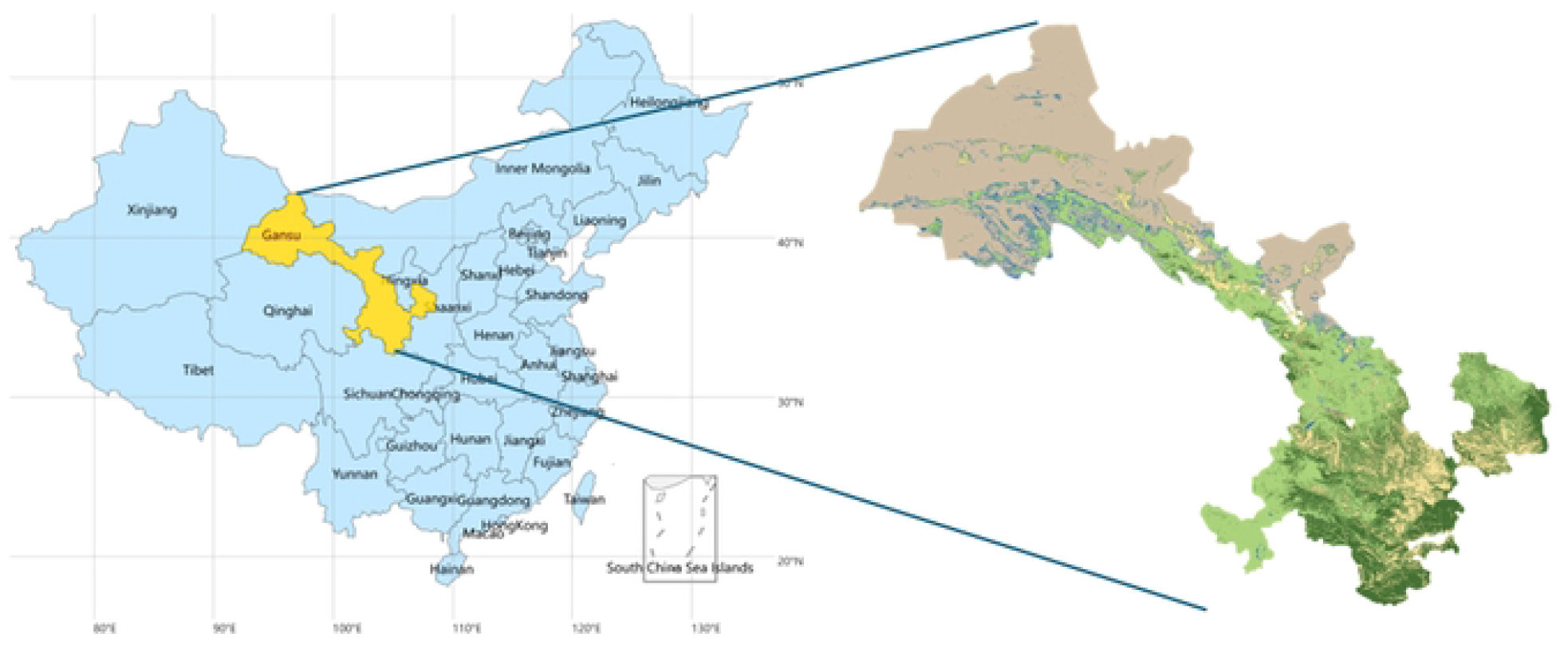

### 2.2 Data sources

The image data used in this study is sourced from the CLCD (China Land Cover Dataset), which is a high-precision, multi-temporal land use/cover dataset released by the Resource and Environmental Science Data Center, Chinese Academy of Sciences. This dataset is primarily constructed based on Landsat remote sensing imagery with a spatial resolution of 30 meters, making it suitable for detailed land use studies at both regional and national scales. The dataset has been classified using a combination of manual visual interpretation and supervised classification with the maximum likelihood method, followed by multiple rounds of correction and validation to ensure the overall data accuracy exceeds 92%, demonstrating high reliability and practicality. The dataset is categorized intom six major land use types, including arable land, forestland, grassland, water bodies, residential areas, and unused land. These are further subdivided into 25 secondary land use types, covering China’s diverse land use patterns and capable of reflecting the spatial and temporal distribution characteristics and dynamic changes of different land types [25].

The CLCD dataset has broad application value, providing critical data support for dynamic land use monitoring and playing a significant role in areas such as ecological environment assessment, resource management, land policy formulation, and sustainable development research. As a high-resolution, widely covered, and highly accurate dataset, the CLCD dataset offers valuable data resources for land use studies in China and globally. It holds considerable research and application potential, particularly in addressing issues such as land resource scarcity, ecological degradation, and rapid urban expansion.

### 2.3 Statistical approach to land use change based on image analysis

This study processes the CLCD dataset using image analysis methods to extract land use data. First, a series of preprocessing steps were applied to the remote sensing imagery in the CLCD dataset, including radiometric correction, atmospheric correction, and geometric correction, in order to eliminate noise and errors in the images, ensuring their accuracy and consistency. After preprocessing, image classification methods were employed to process the land cover data, and the data were reclassified into new land use categories. Through supervised classification of the remote sensing imagery, six major land use types were extracted: arable land, forest, grassland, water bodies, unused land, and built-up land. The reclassification process effectively simplified the diversity of the original data, allowing for clear differentiation between various land cover types and providing clear classification standards for subsequent analysis [26,27].

To visually present the classification results, the study used the Matplotlib library to generate land use type images with color mapping. These images not only clearly indicate the location and distribution of each land use type but also facilitate comparisons between different regions. Each category was assigned a specific color, further enhancing the readability and analytical value of the images. Additionally, the study performed detailed calculations of the areas for different land use types. By considering the spatial resolution of each pixel, the total area for each category was accurately calculated and converted into hectares for easier interpretation and comparison of the distribution of land types. All image processing and data analysis were conducted on a high-performance desktop workstation, equipped with an Intel I9-14900K CPU, an NVIDIA RTX A6000 Ada GPU, and 128GB of memory, providing robust computational support for this study.

### 2.4 Prediction of land use change based on long and short-term memory networks

The process of land use dynamic simulation using the LSTM network is illustrated in Figure 2a, where x_1_, x_2_, …, x_T_ represent the input sample data of the time series, and h_1_, h_2_, …, h_T_ represent the output features of the time series. First, image analysis methods are used to calculate the initial probabilities of land classification for each historical period from the CLCD dataset. The number of neighboring units with the same land use status is then counted, and driving factors are generated. Next, based on the baseline data, random training sample data are generated within the study area. Using long time series land classification data, the LSTM network is employed to calculate the land use development probabilities. Subsequently, global transition probabilities for land use are calculated by introducing disturbance factors, constraint factors, and neighborhood factors, thereby facilitating the dynamic simulation of land use changes.

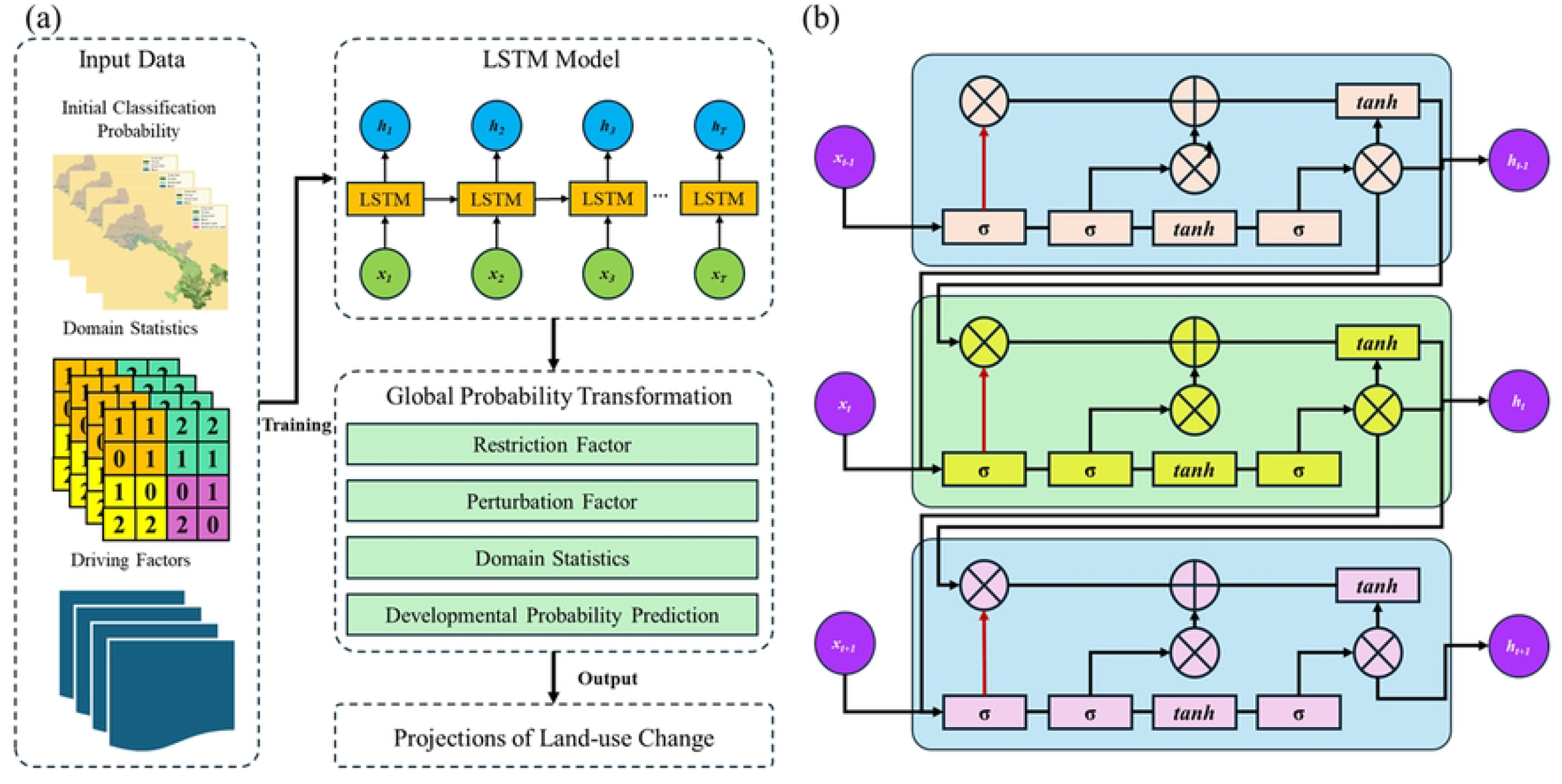

The LSTM network (as shown in Figure 2b), a specialized form of Recurrent Neural Network (RNN), is capable of effectively addressing common issues such as gradient explosion and gradient vanishing during the training process of long time series data. In traditional RNNs, weights are shared across time steps, and gradients are often dominated by those of the more recent time steps, making it difficult for the model to capture long-range dependencies. However, the LSTM network introduces a gating mechanism, which allows the gradients from distant time steps to accumulate along multiple paths, thereby exhibiting superior learning capabilities when handling longer time series data. The basic structure of an LSTM is composed of a series of recurrent units. Compared to ordinary RNNs, LSTM units include input gates, output gates, and forget gates (as shown in Figure 2b), enabling effective filtering and selection of information. In the figure, σ represents the sigmoid activation function, tanh represents the hyperbolic tangent activation function, the red line represents the forget gate, the green line represents the input gate, and the blue line represents the output gate. The x_t_ denotes the input data at the current time step, h_t_ denotes the output feature at the current time step, and ⊕ and ⊕ represent addition and multiplication operations, respectively.

During the model training phase, this study uses land use classification data from 1990 to 2015 and associated driving factors as inputs to the LSTM model. The actual land use data from 2016 serves as a reference, with the model evaluation performed using a cross-entropy loss function. The Adam optimization algorithm is employed to optimize the network parameters. In the prediction phase, the model simulates the development probabilities of land use units for two time periods, 2016-2020 and 2020-2030, providing quantitative predictions for land use dynamic changes.

### 2.5 Space transfer matrix

The land use transition matrix is a matrix used to describe the probability or quantity of transitions between different types of land in a given region over different time periods. It reflects the trends and patterns of conversion between various land use types, providing important decision-making support for land use planning. The formula is as follows:

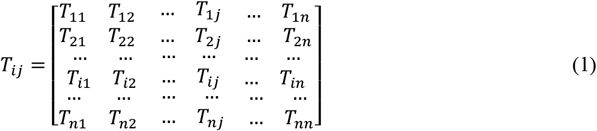

In the formula, *T*_*ij*_ represents the area of land use type i transitioning to land use type j. When i=j, *T*_*ij*_ indicates the area of a land use type that did not undergo any change during the study period. Here, iii and j represent the land use types before and after the transition, respectively, and n denotes the total number of land use types in the study area. As can be seen from the expression, the sum of the rows and columns of the matrix corresponds to the area of a specific land use type at the beginning and end of the study period, respectively.

### 2.6 Analysis of rates of land-use change and dynamics

The land use dynamism model quantitatively reflects the rate of change in the amount of land use within a region and predicts the trend of land use changes. Land use dynamism can be divided into two types: the land use singular dynamism (K) model and the comprehensive land use dynamism (Lc) model. The K model reflects the rate of change in the area of a specific land use type within a given time frame, focusing on analyzing the changes of each land use type. Its calculation method is shown in equation (2). The Lc model describes the overall rate of land use change for the entire region, aiming to reveal the regional differences in land use dynamic changes. Its calculation formula is shown in equation (3).

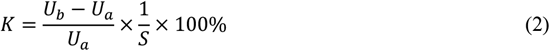

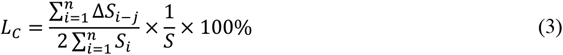

In the formula, *U*_*a*_ and *U*_*b*_ represent the areas (in hectares, hm^2^) of a specific land type at the beginning and end of the study period, respectively; *S*_*i*_ and *ΔS*_*i-j*_ represent the absolute values (in hectares, hm^2^) of the area of land type iii at the beginning of the study period and its converted area; and S denotes the study period. When S is set to a year, the value of *L*_*C*_ represents the annual land use change rate in the study area.

### 2.7 Evaluation of indicators

This study simulates and predicts land use changes in Gansu Province by constructing a time series forecasting model based on LSTM. To evaluate the model’s prediction performance and accuracy, commonly used evaluation metrics were selected. The Root Mean Squared Error (RMSE) is employed to visually reflect the magnitude of the prediction errors, indicating the degree of dispersion of the model’s overall errors. Additionally, to further quantify the model’s prediction accuracy, the coefficient of determination (R^2^) is used as an assessment metric for the goodness of fit. This metric reflects the model’s ability to explain the variation in the dependent variable; the closer the value is to 1, the better the model’s fit. By calculating and comparing these metrics, a comprehensive evaluation of the LSTM model’s performance in land use prediction is conducted, ensuring the reliability and scientific rigor of the results. The specific calculation formulas are as follows:

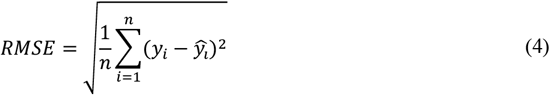

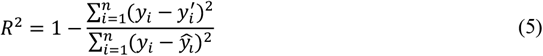

Where *n* denotes the number of objects to be compared; *y*_*i*_ indicates the value of the manual measurement result; 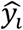 denotes the values of the model; 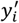 indicates the mean of manual measurement results.

## 3 Results

### 3.1 Analysis of land-use area changes

Based on the statistical data, spatial distribution, and change characteristics of land use types from 1990 to 2020 (Figure 3 and Table 1), various land use types have undergone changes of varying degrees over the 30-year period, with each type experiencing distinct transformation trends. Specifically, the area of built-up land has shown a continuous increase, while unused land has gradually decreased. The trends in arable land, forest, grassland, and water bodies vary. The area of built-up land increased from 34.29 km^2^ in 1990 to 87.32 km^2^ in 2020, with its proportion rising from 0.08% to 0.21%, reflecting the rapid urbanization trend. The area of arable land has fluctuated slightly, increasing from 5,466,325.32 km^2^ in 1990 to 5,834,672 km^2^ in 2000, followed by a slight decrease, reaching 5,511,529 km^2^ in 2020, with its proportion fluctuating from 12.85% to 12.95%.

**Table 1.** Statistical results of land use type changes in Gansu Province in typical years 1990-2020.

| Class | 1990 |  | 2000 |  | 2010 |  | 2020 |  |
| --- | --- | --- | --- | --- | --- | --- | --- | --- |
|  | Area/km<sup>2</sup> | Percentage | Area/km<sup>2</sup> | Percentage | Area/km<sup>2</sup> | Percentage | Area/km<sup>2</sup> | Percentage |
| Cropland | 54663.25 | 12.85% | 58346.72 | 13.71% | 55585.73 | 13.07% | 55115.29 | 12.95% |
| Forest | 34134.41 | 8.02% | 36175.64 | 8.50% | 37629.86 | 8.84% | 40896.56 | 9.61% |
| Grassland | 151229.54 | 35.55% | 147146.73 | 34.59% | 154682.23 | 36.36% | 153035.20 | 35.97% |
| Water | 1758.94 | 0.41% | 164630 | 0.39% | 2084.49 | 0.49% | 2065.22 | 0.49% |
| Unused Land | 183314.73 | 43.09% | 181661.54 | 42.70% | 174794.17 | 41.09% | 173458.33 | 40.77% |
| Construction Land | 342.94 | 0.08% | 466..89 | 0.11% | 667.35 | 0.16% | 873.23 | 0.21% |

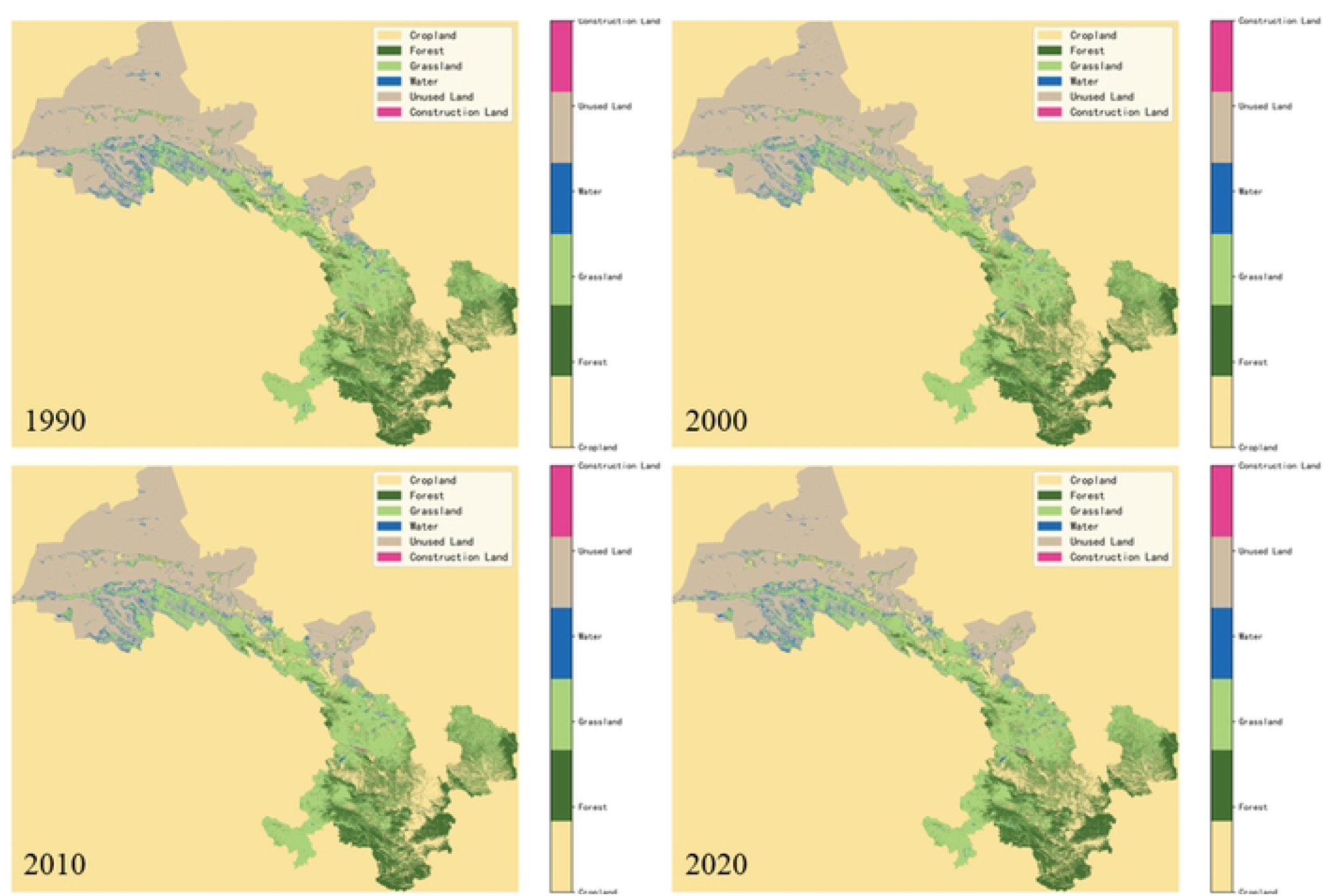

The area of forest land has shown a gradual increase, rising from 3,413,441.16 km^2^ (8.02%) in 1990 to 4,089,656 km^2^ (9.61%) in 2020, reflecting the significant effectiveness of ecological protection and vegetation restoration efforts. The area of grassland has experienced slight fluctuations at different stages but has generally remained stable. It decreased from 15,122,954.2 km^2^ in 1990 to 14,714,673 km^2^ in 2000, then slightly rebounded to 15,303,520 km^2^ in 2020, with its proportion remaining around 35.97%, showing minimal change. The area of water bodies has changed relatively steadily, increasing from 175,894.29 km^2^ (0.41%) in 1990 to a peak of 208,449.5 km^2^ (0.49%) in 2010, before slightly decreasing to 206,522 km^2^ in 2020, maintaining a stable proportion of 0.49%. The area of unused land has gradually decreased, from 18,331,473.7 km^2^ in 1990 to 17,345,833 km^2^ in 2020, with its proportion dropping from 43.09% to 40.77%, indicating a trend of substantial land development and utilization. Regarding the changes in land use types at different stages, the area of built-up land has shown a consistent and rapid increase, reflecting the acceleration of urbanization; forest land has gradually expanded, demonstrating the success of ecological restoration; both arable land and grassland areas have slightly decreased overall, with minimal fluctuation; water bodies first increased and then stabilized; and unused land has exhibited a gradual reduction. From the changes in land use types across periods and the average annual change rates, during the 1990-2000 period, the areas of built-up land, forest, and water bodies increased, while the areas of arable land, grassland, and unused land decreased. The absolute increase in built-up land was the largest, and its average annual change rate was the highest, indicating that the study area was primarily in a period of rapid urbanization. Regional development led to a surge in demand for built-up land, while a large amount of unused land was developed, impacting and partially degrading grassland and arable land resources.

Overall, the spatial distribution and trends of land use changes from 1990 to 2020 indicate that regional urbanization and ecological restoration are the main driving factors behind land use changes. The growth of built-up land and forest areas, as well as the reduction of unused land, show a significant correspondence. These results provide essential data support for further land resource management, ecological protection, and sustainable development planning.

### 3.2 Characterization of the dynamics of land-use change

The single land use dynamics for each land use type from 1990 to 2020 in different time periods were calculated, as shown in Table 2. Overall, the single land use dynamics for each land type exhibited distinct trends of change during the 1990-2020 period. Among them, the change in built-up land was the most dramatic, with the fastest growth rate. The single land use dynamics for built-up land reached 3.9418% per year, showing a continuous acceleration across all periods, especially from 2000 to 2010, when the single land use dynamics peaked at 4.0297% per year, the fastest growth rate. This indicates that during this period, the expansion of built-up land was rapid, and the urbanization process advanced at an accelerated pace.

**Table 2.**
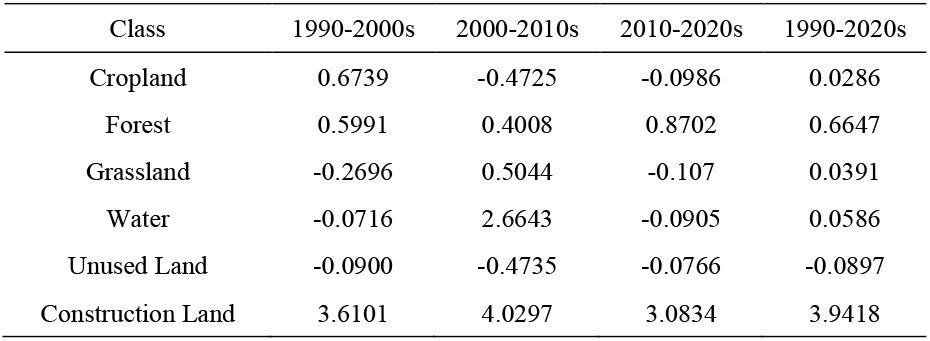
Single-movement attitudes of land use types at typical stages in Gansu Province (unit: %/year)

The overall single land use dynamics for forests was 0.6647% per year, showing a gradual increasing trend, particularly with the highest growth rate between 2010 and 2020, when the single land use dynamics reached 0.8702% per year. This indicates that during this period, the effects of ecological restoration and protection measures were significant, leading to an increase in forest coverage. The overall single land use dynamics for grassland was 0.0391% per year. Although it showed slight overall growth, it fluctuated across different periods, with negative growth observed in both the 1990-2000 and 2010-2020 periods. The single land use dynamics were -0.2696% per year and -0.107% per year, respectively, reflecting the degradation and damage to grassland resources during these phases. For cultivated land, the overall single land use dynamics was 0.0286% per year, with relatively small changes. However, negative growth was observed during the 2000-2010 and 2010-2020 periods, with rates of -0.4725% per year and -0.0986% per year, respectively. This suggests a slight reduction in cultivated land during these periods, possibly due to the encroachment of built-up land. The overall single land use dynamics for water bodies was 0.0586% per year, with relatively stable changes. However, a significant increase in growth rate occurred between 2000 and 2010, with a single land use dynamics of 2.6643% per year, indicating an expansion of water bodies. This was followed by a slight negative growth between 2010 and 2020, with a single land use dynamic of - 0.0905% per year. For unused land, the single land use dynamics was -0.0897% per year, showing a gradual decrease, particularly during the 2000-2010 period, when the single land use dynamics was - 0.4735% per year, marking the most significant decline. This phenomenon suggests that unused land was progressively developed and converted into other land types, especially built-up land.

In summary, between 1990 and 2020, significant differences were observed in the changes of various land use types. Among them, the area of built-up land experienced the fastest growth, with the single land use dynamics showing a continuous acceleration in all periods, reflecting the rapid pace of urbanization. Forests exhibited a steady growth trend, indicating significant success in ecological restoration. The changes in grassland and cultivated land were characterized by periodic decreases, particularly in the 2010-2020 period, where the degradation of both grassland and cultivated land was notably evident. The changes in water bodies were generally small, but there was a significant increase between 2000 and 2010. Unused land continuously decreased, reflecting the gradual development and utilization of land resources.

### 3.3 Analysis of average annual rate of change in land use

The annual average change rates of various land types in Gansu Province from 1990 to 2020, calculated for different time periods, are presented in Table 3. Overall, the annual average change rates of land types from 1990 to 2020 show different trends. Specifically, cultivated land and grassland exhibit periodic fluctuations, while forest and built-up land demonstrate a continuous growth trend. Unused land, on the other hand, shows a gradual decrease over the study period.

**Table 3.** Analysis of the average annual rate of change of land use types at typical stages in Gansu Province (unit: %/year)

| Class | 1990-2000s | 2000-2010s | 2010-2020s | 1990-2020s |
| --- | --- | --- | --- | --- |
| Cropland | 0.0866 | -0.0649 | -0.0111 | 0.0035 |
| Forest | 0.0480 | 0.0342 | 0.0768 | 0.0530 |
| Grassland | -0.0960 | 0.1771 | -0.0387 | 0.0141 |
| Water | -0.0026 | 0.0103 | -0.0005 | 0.0024 |
| Unused Land | -0.0389 | -0.1614 | -0.0314 | -0.0772 |
| Construction Land | 0.0029 | 0.0047 | 0.0048 | 0.0042 |

The annual average change rates of cultivated land, forest, grassland, water bodies, unused land, and built-up land from 1990 to 2020 are as follows. The annual average change rate of cultivated land is generally 0.0035% per year, indicating minor fluctuations and a relatively stable trend. From 1990 to 2000, the area of cultivated land experienced slight growth, with an annual average change rate of 0.0866% per year. However, from 2000 to 2010, the annual average change rate decreased to -0.0649% per year, indicating a reduction in cultivated land area, likely due to the implementation of ecological restoration policies or the expansion of other land types. From 2010 to 2020, the area of cultivated land continued to decrease slightly, with an annual average change rate of -0.0111% per year. Forest area exhibited a stable positive growth trend, with an overall annual average change rate of 0.0530% per year. Between 1990 and 2000, the annual average change rate was 0.0480% per year, showing moderate growth. From 2000 to 2010, the growth rate slightly decreased to 0.0342% per year. However, from 2010 to 2020, forest area increased at an accelerated rate, with an annual average change rate of 0.0768% per year, reflecting the significant effectiveness of ecological restoration measures and a notable increase in forest cover. The annual average change rate of grassland was 0.0141% per year, showing fluctuating changes. From 1990 to 2000, grassland area decreased, with an annual average change rate of -0.0960% per year. Between 2000 and 2010, the grassland area significantly recovered, with an annual average change rate of 0.1771% per year, demonstrating the effectiveness of ecological restoration efforts. However, from 2010 to 2020, grassland area showed a slight negative growth, with an annual average change rate of -0.0387% per year. Water bodies exhibited relatively stable changes, with an overall annual average change rate of 0.0024% per year. From 1990 to 2000, the area of water bodies slightly decreased, with an annual average change rate of -0.0026% per year. From 2000 to 2010, the area of water bodies increased, with an annual average change rate of 0.0103% per year, reflecting the effectiveness of water resource protection and management. From 2010 to 2020, water body changes stabilized, with an annual average change rate of -0.0005% per year. Unused land showed a continuous decreasing trend, with an annual average change rate of -0.0772% per year, the highest decrease among all land types. From 1990 to 2000, the annual average change rate of unused land was -0.0389% per year. From 2000 to 2010, the decrease accelerated, with an annual average change rate of -0.1614% per year. From 2010 to 2020, the reduction rate slightly slowed, with an annual average change rate of -0.0314% per year, reflecting the gradual development and utilization of unused land. The change in built-up land was relatively small but showed a continuous increase, with an overall annual average change rate of 0.0042% per year. Between 1990 and 2000, the annual average change rate was 0.0029% per year. From 2000 to 2010, the growth rate increased to 0.0047% per year. From 2010 to 2020, the growth rate remained stable, with an annual average change rate of 0.0048% per year, indicating the ongoing process of urbanization in Gansu Province and the continuous expansion of built-up land.

In summary, the changes in land use types in Gansu Province from 1990 to 2020 exhibit distinct stage-specific characteristics. Forests and built-up land have shown continuous growth, reflecting the effectiveness of ecological restoration and urbanization development. The changes in grassland and cultivated land have been fluctuating, with reductions observed in certain periods. Unused land has steadily decreased, indicating an increasing intensity of land resource development. The changes in water bodies have been relatively stable, with small fluctuations.

### 3.4 Results of land-use change projections

In this study, we used LSTM to simulate and predict land use from 2016 to 2026, with the results shown in Figure 4. The cultivated land area exhibits fluctuating changes in the time series, without any obvious trend of increase or decrease, remaining relatively stable with periodic fluctuations. The predicted data points are distributed along a slightly upward trend line, suggesting that the cultivated land area may experience a slight increase overall, though with small fluctuations, and the linear fit is moderate. The forest area shows a continuous increasing trend in the time series, with a marked rise in the later periods, reflecting steady growth in forest cover, which may be attributed to ecological protection measures. The predicted data points are highly concentrated along a clearly upward trend line, indicating a stable growth trend for forest area, with a good linear fit. The grassland area demonstrates a U-shaped trend, with a decrease initially followed by an increase, suggesting that grassland resources were degraded or damaged in the early period, and gradually recovered afterward. The predicted data points are distributed along a downward trend line, although recovery is observed in the later periods, the overall trend for grassland area still indicates a decrease, with the linear fit reflecting overall negative growth. The water area exhibits fluctuating changes in the time series, with some ups and downs, showing no significant upward or downward trend. The predicted data points are more dispersed, following a slightly downward trend line, suggesting a slight decrease in water area, though the change is small. The unused land area shows a continuous decreasing trend, with a clear downward slope, reflecting that unused land is gradually being developed and possibly converted into cultivated land or built-up land. The predicted data points are aligned along a clear downward trend line, with a good linear fit, indicating that unused land is steadily decreasing. The built-up land area shows a rapid and continuous increase, with a marked linear rise, reflecting the acceleration of urbanization and infrastructure development. The predicted data points are distributed along a clearly upward trend line, with an excellent linear fit, indicating a stable and rapid growth trend in built-up land area.

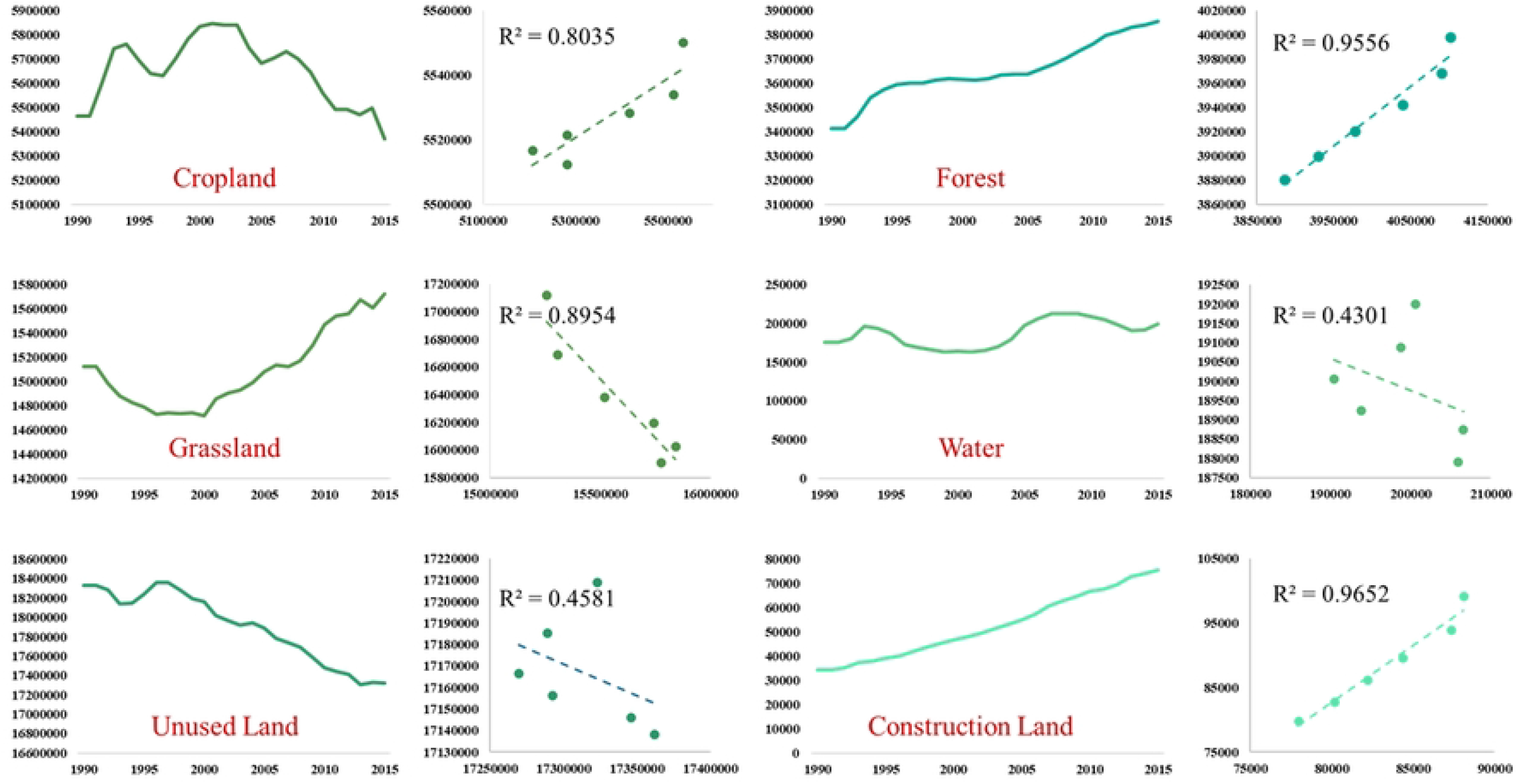

Overall, the rapid growth of built-up land and the continuous reduction of unused land are the most significant land use change trends. The increase in forest area reflects the effectiveness of ecological protection, while the fluctuating changes in grassland and water area highlight the staged characteristics of resource utilization.

### 3.5 Quantitative simulation of land-use change

To achieve a quantitative simulation of future land use changes, our model was used to predict and quantitatively simulate land use changes in Gansu Province for the decade from 2021 to 2030. Table 4 presents the land use type transition matrix of the predicted results. According to the predictions in Table 4, significant spatial transitions occurred between different land use types in Gansu Province during the 2021-2030 period. The change in arable land area was particularly noticeable, with some arable land transitioning to forest, grassland, and unused land, while a substantial portion was converted into water bodies and built-up land. The forest area showed an overall increasing trend, with transitions to grassland and unused land, although some forest land was lost to water bodies and built-up areas. Grassland experienced significant net loss, with a large portion transitioning to other land types, particularly arable land, forest, and unused land, indicating a continued reduction in grassland resources. The change in water area was relatively small and stable, but there was still some conversion of arable land, forest, and grassland into water bodies, with a small portion of water bodies also transitioning to unused land. Unused land showed a substantial reduction during this period, mainly transitioning to arable land, forest, and grassland, with a small amount converting into water bodies. The expansion of built-up land was the most significant trend, absorbing areas of arable land, forest, grassland, and unused land, reflecting the rapid progress of urbanization and infrastructure development. Overall, the land use changes in Gansu Province from 2021 to 2030 show a decrease in unused land, grassland, and arable land, while built-up land, forest, and water bodies are expected to increase. These trends reflect the accelerated development and utilization of land resources, as well as the dual impacts of ecological protection and urbanization.

**Table 4.**
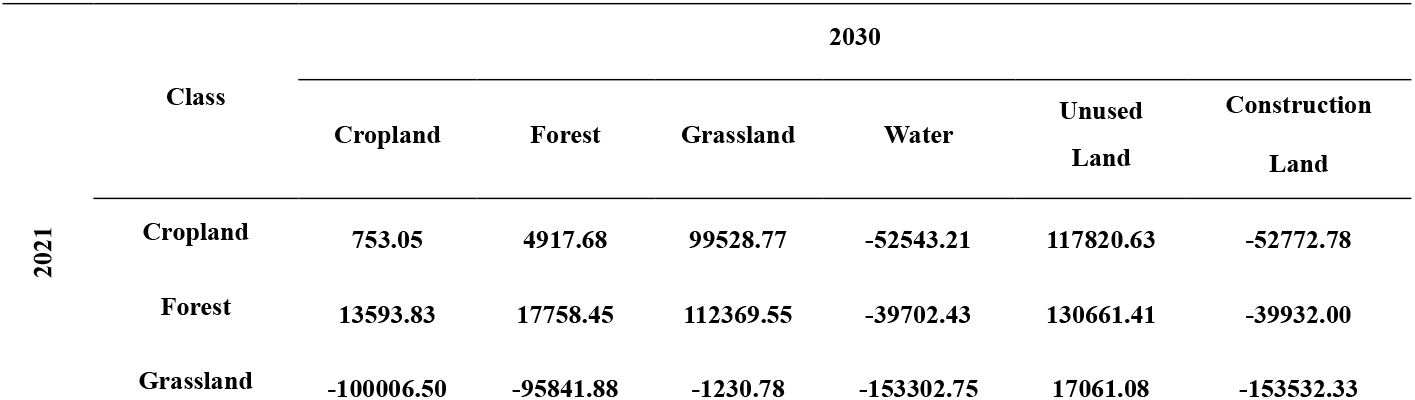
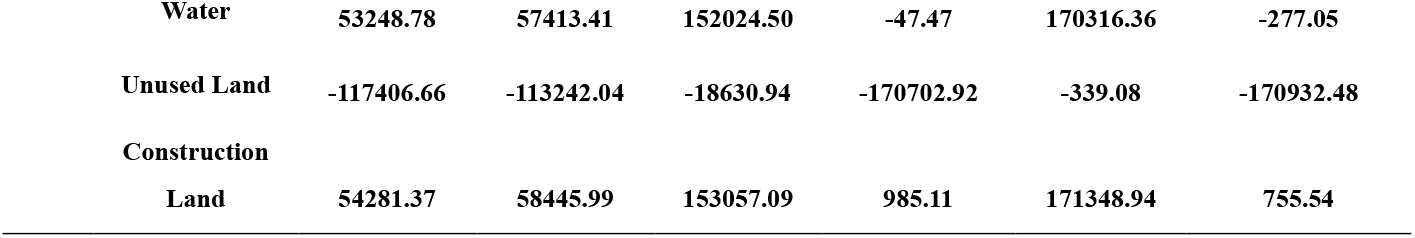
Quantitative simulation analysis of land use type changes in Gansu Province in the next 10 years (unit: km^2^)

## 4 Discussion

### 4.1 Advancement of the methodology

This study takes Gansu Province as a case example and uses land use type data from the past 30 years (1990-2020) to predict future land use changes using the Long Short-Term Memory (LSTM) model. The innovation of this approach lies in its integration of time series analysis and deep learning techniques, enabling it to fully exploit the temporal dependencies and spatial dynamic trends embedded in land use data. As a special type of recurrent neural network (RNN), the LSTM model overcomes the gradient vanishing and exploding problems that are common in traditional RNNs when dealing with long-term dependencies. It can effectively capture the nonlinear patterns of change over long time spans, making it particularly suitable for land use data with strong temporal continuity. In land use change prediction, the LSTM model, through its memory units and forget mechanism, can learn and extract key features and patterns from historical data, allowing for high-precision predictions of future trends. This provides strong scientific support for regional land resource management, planning, and ecological protection [28-30].

Compared to traditional statistical models (e.g., ARIMA) or machine learning-based regression models (e.g., Support Vector Regression, SVR), the LSTM model offers superior nonlinear modeling capabilities and can handle the complex dynamics of land use change. Traditional methods often rely on linear assumptions when processing time series data, while land use change is influenced by both natural factors and human activities, exhibiting high nonlinearity and complexity that traditional methods struggle to model effectively. In contrast, the LSTM model does not require complex preprocessing or feature engineering, as it can directly train on raw time series data and automatically extract regular patterns in land use change over long time spans. Furthermore, the LSTM model can also be combined with other data sources (e.g., climate data, geographical spatial data, human activity data, etc.) to enhance prediction accuracy through multivariate time series analysis, thereby providing technical support for fine-grained land use change modeling [31-33].

### 4.2 Limitations of the methodology

Although the LSTM model demonstrates clear advantages in land use type prediction, it also has certain limitations. First, the LSTM model is highly dependent on the quality and quantity of the data. Land use change data requires high spatial resolution and temporal completeness, but in practical applications, obtaining long-term continuous high-quality land use data can be challenging. Incomplete or noisy data may adversely affect the model’s training performance and prediction accuracy [34]. Furthermore, due to the high computational complexity of the LSTM model, the training process can be time-consuming and demanding on hardware resources. In this study, high-performance computing systems (e.g., workstations equipped with GPUs) were necessary to support the training of large-scale time series data. This may pose a challenge for researchers with limited resources [35].

Secondly, the LSTM model, when applied to land use type changes, cannot directly incorporate spatial feature information. Land use data not only exhibit temporal dynamics but also display significant spatial heterogeneity and spatial autocorrelation. While the LSTM model is primarily designed to model the temporal dimension, it struggles to capture the spatial patterns of land use type changes. Therefore, relying solely on the LSTM model for prediction may limit the accuracy of spatial pattern change predictions. Future research could combine the LSTM model with spatial analysis methods, such as Geographic Information Systems (GIS), remote sensing technologies, or spatially explicit deep learning models (e.g., Convolutional Neural Networks, CNN), to construct spatiotemporal joint prediction models, thus improving the reliability and applicability of the predictions. Additionally, the poor interpretability of the LSTM model is another important limitation in practical applications. While the LSTM model can capture nonlinear features in time series data effectively, its internal structure is complex, making it difficult to intuitively explain the driving factors behind each prediction. In land use change research, scientific decision-making and policy formulation require not only understanding the prediction results but also clearly identifying the key factors influencing land use change and their underlying mechanisms. Therefore, model result interpretation and visualization are key directions for future research. Combining methods like sensitivity analysis and variable contribution assessment could provide clearer explanations for the prediction outcomes [36, 37].

In summary, this study utilizes the LSTM model to predict land use types in Gansu Province, characterized by its advanced methodology and strong applicability, providing scientific support for land resource management and future planning. However, it also has certain limitations, such as high data dependency, difficulty in incorporating spatial features, and poor interpretability. Future research could enhance the LSTM-based land use change prediction method by integrating more multi-source data, constructing spatiotemporal joint models, and improving result interpretability, thereby promoting its practical application in land management and regional sustainable development.

## 5 Conclusion

This study, based on land use type data from Gansu Province between 1990 and 2020, constructs a high-precision time series model to predict land use changes over the next decade. The results show that the dynamic changes in different land use types exhibit distinct phase characteristics, reflecting the combined impact of ecological restoration and human activities. The changes in forest land and built-up land are the most notable. Forest area continues to increase throughout the prediction period, with the growth rate accelerating particularly between 2020 and 2030, reflecting the effectiveness of ecological protection policies and vegetation restoration measures. Built-up land expands rapidly, absorbing substantial areas from cropland, grassland, and unused land, indicating the accelerated pace of urbanization and infrastructure development. The changes in grassland are complex and volatile. Although there is some recovery in certain stages, the overall trend is a decline, suggesting the dual pressure of ecological degradation and development. Cropland area remains generally stable, but decreases slightly in certain stages, possibly due to the encroachment of built-up land. The reduction of unused land is the most significant, with it continuously being developed and mainly converted into cropland and built-up land. The water area remains stable overall, with fluctuations but no significant increase or decrease, likely related to natural variations and management measures.

The study demonstrates that the LSTM model has significant advantages in capturing land use change trends, providing scientific support for regional land resource management and policy formulation. However, the model has limitations in capturing spatial features of land use changes. Future research could integrate spatial explicit models and GIS technology to enhance simulation capabilities. Additionally, the interpretability of the model’s predictions is low and needs to be improved through sensitivity analysis or driver mechanism models. These conclusions provide valuable references for sustainable land use and regional development and offer directions for future research.

## CRediT authorship contribution statement

Shiqi Zhang: Conceptualization, Investigation, Methodology, Visualization, Writing – original draft. Chuhui Cao: Resources, Funding acquisition, Supervision.

## Acknowledgments

This work was partially supported by Social Science Foundation of Beijing Union University.

## References

1. Wu Fei, Zhang Lei, Chen Xiaoming, Li Wei, Liu Qing. *Time-Series Analysis of Cropland Changes Using ARIMA*. Land Use Policy, 2018.

2. Li Cheng, Wang Jie, Zhou Hua, Zhang Ying, Liu Fang. *Urban Expansion Prediction Using LSTM Networks*. Remote Sensing of Environment, 2020.

3. Zhang Yong, Sun Ming, Liu Haoran, Wang Qiang, Chen Xiang. *Predicting Forest Cover Changes with Deep Learning*. Applied Geography, 2021.

4. Wang Jun, Zhao Liang, Li Mei, Gao Feng. *Land Use Changes and Their Impacts on Ecosystem Services in Gansu Province*. Journal of Environmental Management, 2019.

5. Zhang Hongwei, Liu Xiaoming. *Forest Restoration Policies and Their Impacts in China*. Environmental Science & Policy, 2021.

6. Liu Jingyun, He Qing, Zhang Yuan, Li Xiaoxia. *Spatio-Temporal Dynamics of Land Use in Northwest China*. Geographical Research, 2014.

7. Huang Qinghua, Zhang Mingyi, Sun Honglei. *Land Use Change in China and Its Driving Forces*. Land Use Policy, 2019.

8. Long Hualou, Liu Yansui, Wu Xiaofang, Zhang Yu. *Analysis of Land Use Transition in China*. Habitat International, 2012.

9. Verburg Peter H., Overmars Koen P., Witte Nadia. *Modeling Land Use Change and Its Environmental Impact*. Journal of Environmental Management, 2006.

10. Yang Xiaohong, Huang Huimin. *Remote Sensing and GIS-Based Land Use Analysis in Northwestern China*. Environmental Monitoring and Assessment, 2020.

11. Chen Xiang, Zhang Li, Wang Hao, Zhang Qing. *Machine Learning Models for Land Use Classification*. ISPRS Journal of Photogrammetry and Remote Sensing, 2019.

12. Wu Jianhua, Zhao Rong, Chen Jie, Gao Zhiyong. *Impacts of Land Use and Climate Change on Ecosystems*. Global Environmental Change, 2017.

13. Song Weili, Zhang Huifeng, Liu Qian, Zhao Fang. *Evaluating Urbanization Impact on Agricultural Land*. Environmental Science and Policy, 2021.

14. Gao Bin, Xu Lei, Wang Xiaolin. *Remote Sensing Data Fusion for Land Use Monitoring*. Remote Sensing Letters, 2020.

15. Luo Xianjun, Chen Jianfeng, Hu Xiangdong. *Spatio-Temporal Analysis of Land Degradation in Arid Regions*. Journal of Arid Environments, 2019.

16. Jiang Lijun, Zhang Zhihong. *Application of LSTM in Predicting Land Use Dynamics*. Computers and Geosciences, 2020.

17. Bai Yunfei, Zhao Wei, Liu Qinghua, Zhang Cheng. *Impacts of Land Use Change on Soil Erosion in the Loess Plateau*. Geomorphology, 2018.

18. Zhao Wei, Zhang Ronghua, Liu Fang. *The Role of Policies in Land Use Transition in China*. Land Degradation & Development, 2016.

19. Fischer Günther, Velthuizen Harrij, Nachtergaele Freddy. *Global Agro-Ecological Zones: Modeling Land Use and Climate Change*. Agricultural Systems, 2010.

20. Fang Hui, Li Pengfei, Wu Xing, Zhang Yan. *Temporal and Spatial Patterns of Land Use Changes in Arid Regions*. Remote Sensing, 2019.

21. Turner Billie Lee III, Lambin Eric F., Reenberg Anette. *The Emergence of Land Change Science for Global Environmental Change*. PNAS, 2007.

22. Zhao Shuang, Li Qing, Zhang Xiang, Wang Rui. *Urban Land Expansion and Its Driving Forces in China*. Ecological Indicators, 2018.

23. Yu Xiaowei, Zhang Feng, Li Hongwei. *Dynamic Simulation of Land Use with Integrated Models*. Journal of Environmental Informatics, 2020.

24. Zhu Qian, Liu Wen, Zhang Huan, Zhou Xing. *Integrating LSTM and Remote Sensing Data for Land Change Prediction*. Sustainability, 2021.

25. Zhang Sheng, Ma Ling, Yang Yan, Zhao Jun. *Driving Forces of Land Use Changes in Gansu Province*. Journal of Cleaner Production, 2020.

26. Ren Jianhua, Wang Yan, Li Tao. *Assessment of Land Use Change and Its Impacts on Water Resources*. Water, 2018.

27. Yang Jie, Liu Xiaobo, Wang Hua, Zhang Lei. *Impacts of Human Activities on Grassland Degradation in China*. Journal of Environmental Management, 2019.

28. Zhang Chao, Li Rong, Wang Yifan. *A Spatio-Temporal Model for Predicting Land Use Dynamics*. Land Use Policy, 2020.

29. Batty Michael. *Cities and Complexity: Understanding Land Use Dynamics*. Environment and Planning B, 2007.

30. Zhang Xiaoyu, Chen Jie, Wang Yong. *Combining LSTM and GIS for Spatial-Temporal Land Use Prediction*. Applied Geography, 2021.

31. Zhu Xiangdong, Wu Jing, Zhang Yanan. *Urbanization and Land Use Change in China*. Land, 2016.

32. Liu Yansui, Zhang Zhaoyu, Wu Tian. *The Role of Policies in Shaping Land Use Trends*. Journal of Geographical Sciences, 2019.

33. Liu Chao, Sun Huan, Zhang Ming. *Effects of Land Use Transition on Ecosystem Services*. Ecological Indicators, 2018.

34. Wu Hongbo, Zhao Xia, Li Jian. *Implications of Urbanization on Land Use Planning*. Urban Studies, 2020.

35. Zhou Jianfeng, Liu Zhichao, Wang Han. *Integrating Remote Sensing and LSTM for Regional Land Use Dynamics*. Land Degradation & Development, 2021.

36. Tang Liang, Wu Min, Zhang Qingyuan. *Multi-Source Data Integration for Land Use Monitoring*. Journal of Remote Sensing, 2020.

37. Li Min, Zhao Yu, Zhang Fang. *Forecasting Agricultural Land Use Using LSTM*. Agriculture, 2021.

